# APICURON: a database to credit and acknowledge the work of biocurators

**DOI:** 10.1101/2021.02.03.429425

**Authors:** András Hatos, Federica Quaglia, Damiano Piovesan, Silvio C.E. Tosatto

## Abstract

APICURON is an open and freely accessible resource that tracks and credits the work of biocurators across multiple participating knowledgebases. Biocuration is essential to extract knowledge from research data and make it available in a structured and standardized way to the scientific community. However, processing biological data - mainly from literature - requires a huge effort which is difficult to quantify and acknowledge. APICURON collects biocuration events from third party resources and aggregates this information, spotlighting biocurator contributions. APICURON promotes biocurator engagement implementing gamification concepts like badges, medals and leaderboards and at the same time provides a monitoring service for registered resources and for biocurators themselves. APICURON adopts a data model that is flexible enough to represent and track the majority of biocuration activities. Biocurators are identified through their ORCID. The definition of curation events, scoring systems and rules for assigning badges and medals are resource-specific and easily customizable. Registered resources can transfer curation activities on the fly through a secure and robust Application Programming Interface (API). Here we show how simple and effective it is to connect a resource to APICURON describing the DisProt database of intrinsically disordered proteins as a use case. We believe APICURON will provide biological knowledgebases with a service to recognize and credit the effort of their biocurators, monitor their activity and promote curators engagement.

Database URL: https://apicuron.org

## Introduction

Biocuration plays a key role in making research data available to the scientific community in a structured and standardized way. Managing biological data in the biocuration context requires ensuring their accuracy, accessibility and reusability (1,2). Biocurators deal with unstructured data, mainly derived from publications, that need to be collected, verified, standardized and integrated into dedicated databases (1,2). Even though in principle expert curation scales well with the growth of scientific literature, it is highly demanding, requiring the allocation of dedicated resources and time (3,4).

Despite its importance, the contribution and effort of biocurators is extremely difficult to acknowledge and quantify (5). This is relevant for all categories of curators irrespective of they being professional curators, volunteer curators or just experts which occasionally provides feedback on existing annotation records. The European life science infrastructure for biological information (ELIXIR, https://elixir-europe.org/), the International Society for Biocuration (ISB, https://www.biocuration.org) (6) and the Global Organization for Bioinformatics Learning, Education and Training (GOBLET, https://www.mygoblet.org) (7,8) recognized the need to ensure that the role of biocurator is visible and valued (5). According to a global survey they found that biocurators have diverse job titles, are highly skilled, perform a variety of activities and use a wide range of tools and resources. But the majority of biocurators are unaware of this as a career, or simply think that biocuration is not the main aspect of their job (5).

Professional curators are aware of what their role and duties are, as well as their value to the community, for volunteer or occasional curators this is less clear. For both professional and community curators crediting and quantifying their efforts is problematic and a simple way to engage volunteers does not exist.

An established strategy for crediting curators is to offer co-authorship in the article describing the corresponding curated database. However, publication is not always guaranteed or can just be occasional, at the best every two years as stated by Nucleic Acid Research database issue rules (9) (https://academic.oup.com/nar/pages/ms_prep_database).

Moreover, the motivation of community biocurators has been observed to significantly increase and decrease immediately before- and after-publication respectively (10), whereas a consistent effort is required to regularly maintain and extend the data collection. In addition, curation contributions can come from experts in the field who are not entering a database publication, for example those authors that simply provide feedback on annotations referring to their publications. All these cases are never acknowledged and often biocurators work behind the scenes in projects where biocuration is needed, e.g. construction of specific datasets (11–13).

Several efforts have been made during the last decade to credit the work of biocurators in manually curated resources. The Pfam database of protein families (http://pfam.xfam.org/) relies on an ORCID-based system. Biocurators can link their contributions to their ORCID profile claiming Pfam curated entries (14) and their contribution is discoverable by means of the European Bioinformatics Institute EBI-Search system (15). The Reactome Knowledgebase (16) (https://reactome.org) also uses ORCID to credit curation of biochemical pathways. The mission of the Open Researcher & Contributor ID (ORCID, https://orcid.org) is to provide a registry of persistent unique identifiers for researchers and scholars (17). ORCID allows integration of ‘Works’ which are publications, data sets, conference presentations, etc. in the user profile page. However, ORCID is not designed to directly manage biocuration activities and it does not provide a system to aggregate and weigh this type of information. Pfam does not provide a statistics page dedicated to curators. To enumerate or quantify curation contributions it is necessary to download and manually process all database records. Reactome provides curator level statistics, but it is necessary to explicitly search for the curator identifier or name.

Some databases already provide curation statistics and leaderboards. For example the Clinical Interpretation of Variants in Cancer database (CIViC, https://civicdb.org/home) (18) implemented a sophisticated system which takes into consideration biocuration roles. The “*curators*” submit new pieces of evidence while “*editors*” validate new annotations by accepting or rejecting the curators’ work. This is similar to the “curator” and “reviewer” system implemented in DisProt (19).

CIViC also implements some of the core concepts of gamification. Gamification consists of providing motivational affordances and “gameful” experiences in non-game related settings. It is meant to support user engagement and positive behavioural patterns that lead to an overall increase in the quality and productivity of users’ activities (20,21). A key concept of gamification consists of providing users with badges and scores associated with their achievements, giving rise to a positive feedback mechanism that keeps users engaged and motivated and increases their competitiveness (20,21). Gamification concepts apply to all ages and types of people. CIViC includes dedicated profile pages for all biocurators. Contributions are clearly tracked and biocurators accumulate activity scores and badges and they are ranked in a dedicated leaderboard page (18).

However, these tracked contributions pertain only to the biocuration activities provided in CIViC and no similar tracking system is currently available for other manually curated resources. Biocurators working for different organizations or databases do not have a shared place to record their achievements. Curation activities in different resources can be extremely different and can be tracked at different levels of granularity. Moreover, it is extremely difficult to quantify the curation effort for all possible activities.

Here we provide a comprehensive description of the implementation of APICURON and the gamification concepts used. APICURON aims at providing biological databases and organizations with a real-time automatic tracking system of biocuration activities. APICURON, regardless of the field of interest, is a unified resource able to properly attribute and recognize the work of biocurators. APICURON implements badges, medals and leaderboards that promote engagement and allow to evaluate objectively the volume and quality of the contributions. The achievements and scoring systems are defined by the member databases themselves as different resources need to capture different types of activities and want to value biocuration actions differently. APICURON provides support to member databases in the registration process for the definition of the achievements and to identify the best granularity of the curation activities to be tracked. APICURON cross-references the annotation source and activity records are uniquely identified. Every change in the source database, including changes in existing records, can be (re)submitted at any time. APICURON is agnostic about the choices made by databases about how to track curation activity and it is robust to different definitions of activities and scores. APICURON can also be considered stateless since whenever activity records are submitted (or resubmitted) all achievements are recalculated and updated.

The web interface provides both a resource- and curator-centric view including personal pages summarizing biocurators achievements for all resources they work for. Finally we explain in detail how a partner resource can connect to APICURON and submit biocuration events by describing the integration of the DisProt database of intrinsically disordered proteins (IDPs) (19) as a use case.

## Database structure and implementation

APICURON is a web server accessible through a public REStful API or through a user friendly web interface built on top of the same API. It can be used as an external tracker by third party resources and by the biocurators themselves to demonstrate and value their contributions. APICURON is easy to connect with third party databases allowing small emerging databases to adopt best practices and save the cost of implementing the technology for tracking curators work and provides profile pages which showcase biocurators contributions across multiple resources in a single view. In the following paragraphs we describe the philosophy and technical solutions adopted by APICURON to track curation activities, implement gamification experience and manage data transfer (transactions) with registered resources and organizations. An example about how a database can connect to APICURON is provided in the “DisProt use case” section.

### Biocuration activity records

The main feature of APICURON is the tracking of biocuration activities or events. A biocuration event is identified by four fields, *entity*, *activity*, *agent* and *timestamp*, fulfilling the recommendations provided in (22). In APICURON the *entity* corresponds to the curated data object, which is possibly identified by using the identifiers.org resolving system (23). The *activity* is the actual curation activity, such as “entry creation” or “entry invalidation”. Although different databases can define different activities, those are required to be mappable to Provenance (PROV) Ontology terms as defined by the World Wide Web Consortium (W3C) Provenance Working Group (24). PROV terms, *generation*^1^, *revision*^2^ and *invalidation*^3^, provide a way to define the life cycle of an object. In APICURON the *agent* is the biocurator that has performed the biocuration activity and it is identified by the ORCID (17) in order to avoid duplicates or wrong attributions. Finally, the timestamp provides the date and time of the biocuration activity.

The granularity of curation activity records is connected with the ability of a resource to capture curation activity on parts of the curated object or entry. What is a curated object and how many parts it has is not a concern of APICURON. The only constraint set by APICURON is the encapsulation of each activity under PROV categories. A possible standardization of more specific subcategories, eg. with a dedicated ontology or controlled vocabulary, once developed, will be adopted effortlessly.

### Achievements and ranking

APICURON implements gamification by providing badges, medals and leaderboards. They are meant to encourage participation and commitment and promote engagement. *Badges* are based on an absolute count of the activities and represent biocurators milestones or achievements. They are not competitive as they are independent from those of other curators, e.g. “First paper curated” or “Senior biocurator”. On the other end *medals* are competitive as based on the relative performance of each biocurator and depend on the ranking, eg. “Best biocurator 2020” or “Top 10 biocurator 2020”. Since biocuration records have a timestamp, medals can be defined on a time frame, e.g. “Best biocurator 2021”. In order to provide flexible and personalized achievements, each partner resource or database can define different rules to assign badges and medals. Another gamification concept is the leaderboard which represents the global ranking across all players. Different databases might track different types of activities or might want to evaluate different activities in different ways. In APICURON different databases are treated as independent entities and rankings and achievements are never combined. Leaderboards are calculated based on the amount of curation activities and the score associated with each type of activity.

The scoring system allows databases to weigh different curation events. The activity score can be intended as proportional to the amount of time spent by a curator, but it can also be interpreted as the importance of an activity from the point of view of a member database. Some activities can be prioritized by increasing their score and therefore orient curators’ focus.

### Web server

The APICURON workflow is described in Figure 1. APICURON is a web server which exposes a RESTful API. Some endpoints are private and serve for the submission of curation activities from member databases. Public endpoints provide achievements and rankings for third party clients and are also used by the APICURON website. Biocurators activity can be submitted at any time by registered resources. Whenever new data is submitted, biocurator achievements and ranking are calculated (or recalculated) and made available to the web interface (website) or third party services, including the registered resources themselves and other services like ORCID. The list of endpoints and their functionality are provided in Table 1.

This strategy allows the member databases to delegate completely the tracking management and gamification (storage, versioning, achievement and ranking calculation) to APICURON. All data are cross-referenced and stored permanently and the design is flexible enough to accommodate different types of biocuration activities. APICURON can manage different rewording strategies, never mixing records coming from different resources. Member (or client) resources that decide to connect to APICURON are requested to register a new account, which has no cost. At registration time they have to provide the full list of all possible biocuration activities and the corresponding scores, which are used to calculate medals and the leaderboard. In addition they are requested to provide achievements definitions, i.e. the rules for assigning badges and medals.

Submitted data are upserted in the database using the *entity*, *activity*, *agent*, *timestamp* quadruplet as identifier, so that existing records with the same identifier can be overwritten. APICURON does not create an internal identifier so it is the responsibility of the registered resource to decide whether to preserve or overwrite existing records.

The server is implemented using Node.js framework (https://nodejs.org) and stores data in MongoDB database (https://www.mongodb.com). All communications travel through the HTTPS protocol and private communications are protected by an authentication system with JSON Web Token (JWT) credentials.

**Table 1.** APICURON API endpoints. All endpoints are to be prefixed with the https://apicuron.org/api/ domain fragment. Private endpoints can be used only by registered databases to submit biocuration activities. All responses are objects in JSON format. Full documentation is available on the APICURON website.

| Endpoint | Accessibility | Type | Purpose |
| --- | --- | --- | --- |
| submit_description | private | post | Store information about a member resource (list of activity terms and scores, badges, medals, resource description) |
| submit_activity | private | post | Store biocuration events of a member resource |
| delete_activity | private | get | Delete biocuration events of a member resource. Parameters: start/end date, ORCID, etc. |
| reports | public | get | Return biocuration activities. Parameters: resource URI, ORCID, etc. |
| achievements | public | get | Return biocurators achievements (badges, medals). Parameters: resource URI, ORCID, etc. |
| leaderboards | public | get | Return biocurators ranking. Parameters: resource URI, ORCID, etc. |

**Figure 1.**
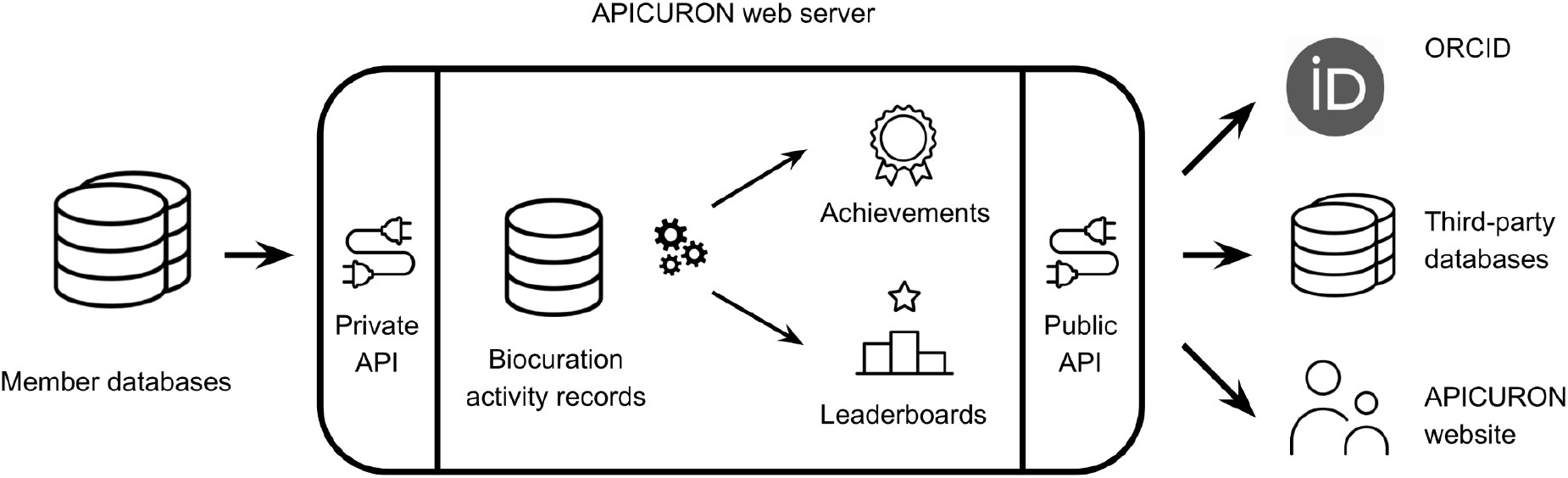
APICURON workflow. Registered resources (Member databases) submit biocuration events to the APICURON web server. APICURON stores biocuration activities and calculates achievements (medals, badges) and leaderboards on the fly. Results are served through a public API. APICURON can be cross-referenced by ORCID. Third-party databases can use APICURON to monitor biocurators work and human users can visit the APICURON website to see biocurators achievements and ranking.

### Website

APICURON both promotes biocurators engagement and provides a monitoring service for registered resources (member databases). The APICURON web interface provides two different perspectives to highlight biocuration work, one focusing on biocurators themselves (personal page) and another on biocurators as contributors of a resource (database page). The curator personal page provides the complete history of the activities, badges and medals and some biographical information as provided by the ORCID public API. A simple statistics provides the amount of activity aggregated at the level of the corresponding PROV category. Medals and badges are provided as graphical elements and colored based on type and importance. Only earned medals are displayed, whereas all badges are always visible but only those achieved appear as activated (highlighted). Curators can decide to hide completely their profile page and all activities connected with their ORCID by simply logging-in using their ORCID credentials and toggling off public visibility of their data.

The database page allows to visualize information for a specific database including the full list of available medals and badges, the leaderboard and the full history of biocuration activities. By clicking on medals and badges is it possible to update the leaderboard table and see the list biocurators who earned a given medal or badge.

## DisProt use case

DisProt is a manually curated database of intrinsically disordered proteins (IDPs) and regions (IDRs) (19). DisProt annotations are provided by a community of expert biocurators who are rewarded with co-authorship in the main database publication every three years. All annotations are checked and validated by a small team of reviewers. DisProt tracks curation work and provides a rudimentary leaderboard, however proper recognition is missing. Here we showcase how DisProt can value the biocurators’ effort qualitatively and quantitatively with APICURON.

A DisProt annotation is a piece of evidence of an IDR which must include an experimental detection method, the reference to the corresponding publication and sequence boundaries of the disordered region. In DisProt each evidence has a stable region identifier, e.g. DP00086r067, an IDR evidence of the human tumor suppressor p53.

### DisProt achievements

DisProt implements versioning in order to track all IDRs changes between different releases. Each time a biocurator fills-in new data in the DisProt curation interface, a copy of the previous version is stored in a history collection and the new object replaces the old one in the active collection.

In DisProt annotation events which are submitted to APICURON are defined by comparing the current version of an IDR with the previous version. For each IDR DisProt compares all fields one-by-one across the version history and maps the differences to a controlled vocabulary of activity terms. DisProt performs this calculation and submits curation activities to APICURON every day in order to keep biocurators engaged. Some activity terms and the corresponding PROV categories are reported in Table 2.

While activities are directly reported by the client database, medals and badges are calculated by APICURON based on the rules defined by DisProt at registration time. Different databases can define different types of activities and different rules for badges and medals. In Table 2, 3 and 4 are reported some examples for DisProt. The “Newbie curator” is achieved when a biocurator totals at least ten activities of any type. To calculate medals for each biocurator APICURON multiplies the number of activities by the corresponding score and ranks the biocurators. Time-dependent medals, e.g. “Best biocurator 2017”, are calculated by filtering activities by date before generating the ranking.

**Table 2.** Example of activity terms as defined in DisProt. Category corresponds to the PROV classes accepted by APICURON. Term is the internal identifier of an activity while Score is used to calculate the ranking and assign medals.

| Category | Term | Score |
| --- | --- | --- |
| generation | region_reference_first_generation | 50 |
| generation | region_first_generation | 20 |
| generation | region_statement_generation | 5 |
| generation | region_crossref_generation | 5 |
| revision | region_validation_revision | 20 |
| revision | region_statement_revision | 5 |
| revision | region_crossref_revision | 5 |
| invalidation | region_invalidation | 20 |

**Table 3.** Example of badges as defined in DisProt. Group is an identifier for those badges evaluated on the same set of activity terms. (*) “All activity terms” indicates that the corresponding badges are evaluated considering all possible types of biocuration activities.

| Group | Name | Count | Activity terms |
| --- | --- | --- | --- |
| 1 | Newbie curator | 10 | all activity terms <sup>a</sup> |
| 1 | Experienced curator | 100 | all activity terms <sup>a</sup> |
| 1 | Advanced curator | 500 | all activity terms <sup>a</sup> |
| 1 | Senior curator | 1000 | all activity terms <sup>a</sup> |
| 1 | Curator ace | 5000 | all activity terms <sup>a</sup> |
| 4 | Novel publication in database | 1 | region_reference_first |
| 4 | Bibliophile | 10 | region_reference_first |
| 4 | Librarian | 50 | region_reference_first |
| 4 | Book worm | 100 | region_reference_first |
| 4 | Book addicted | 1000 | region_reference_first |

**Table 4.** Example of medals as defined in DisProt. Group is an identifier for those medals evaluated on the same time frame (From/To). Ranking indicates the curator ranking threshold to achieve a medal. (*) “All activity terms” indicates that the corresponding medals are evaluated considering all possible types of biocuration activities.

| Group | Name | Ranking | From | To |
| --- | --- | --- | --- | --- |
| 1 | All time best biocurator | 1 |  |  |
| 2 | Best biocurator 2016 | 1 | 1/1/2016 | 31/12/2016 |
| 3 | Best biocurator 2017 | 1 | 1/1/2017 | 31/12/2017 |
| 1 | All time top 10 biocurator | 10 |  |  |
| 2 | Top 10 biocurator 2016 | 10 | 1/1/2016 | 31/12/2016 |
| 3 | Top 10 biocurator 2017 | 10 | 1/1/2017 | 31/12/2017 |
| 1 | All time top 25 biocurator | 25 |  |  |
| 2 | Top 25 biocurator 2016 | 25 | 1/1/2016 | 31/12/2016 |
| 3 | Top 25 biocurator 2017 | 25 | 1/1/2017 | 31/12/2017 |

### How to become a member database

We welcome all manually curated resources or biocuration organizations, especially those relying on community curation, to become APICURON partners. Becoming an APICURON partner is free of charge. To connect to APICURON a resource delegate needs to contact the APICURON team to plan their involvement and if necessary receive support for the definition of the biocuration activities, associated scores, badges and medals. Each registered resource can directly submit data through the APICURON API after receiving credentials. By default, once a manually curated resource becomes a partner database, all biocurators profiles and curation activities are public. However, biocurators can login with their ORCID credentials and toggle off public visibility.

We encourage the submission through the API but we are still open to include small or brand new resources that do not have enough resources to implement a web server able to push data. In that case APICURON developers will take care of collecting (pulling) curation activities manually directly from the connecting resource.

APICURON serves as an external tracker of biocuration activities of each one of its partner resource, playing a crucial role not only in well-known and established databases but also in smaller, newer manually curated resources and, behind a firewall, also for non-academic institutions and research companies where biocuration is required with the aim of properly crediting the work of biocurators, raising awareness to the crucial role played by biocuration in scientific research and promoting good practices.

The APICURON team can be contacted at.

## Conclusion and future work

Here we presented APICURON, a resource to track and acknowledge the work of biocurators. In order to engage biocurators APICURON implements gamification concepts by introducing achievements and leaderboards. APICURON stores biocuration events as provided by registered resources and calculates achievements and ranking based on custom rules and scoring schemas. Biocuration records can be provided at any time and achievements get updated on the fly. Each database defines scores and rules and has its own leaderboard.

The APICURON web interface provides both a biocurator- and resource-centric viewpoint. Biocurators are identified by their ORCID so that they can report APICURON achievements in their *curriculum vitae*. A robust API allows the integration of APICURON results into third-party resources and possibly, in the future, directly into ORCID.

APICURON already includes the biocuration activities as provided by DisProt. We hope that a wider use of APICURON will increase awareness of the crucial role played by biocurators in making life science data available in a well-structured way. We also hope that the knowledge and skills offered by biocurators will be better recognized in career paths as strongly recommended by ELIXIR, ISB and GOBLET organizations. We encourage curation databases, including small and newer resources, to connect to APICURON therefore adopting best practices for keeping biocurators, especially community ones, motivated and promoting their visibility. In order to harmonize as much as possible the quantification of the curation efforts between different resources the APICURON consortium promotes an open discussion with the wider community of curators.

## Acknowledgments

This project has received funding from the European Union’s Horizon 2020 research and innovation programme under grant agreement No 778247 and No 823886. Funding from the Italian Ministry of University and Research (MIUR) Progetti di Ricerca di Interesse Nazionale, PRIN (Grant No. 2017483NH8). This work was funded by ELIXIR, the research infrastructure for life-science data-ELIXIR Data Platform.

## Footnotes

1 https://www.w3.org/TR/prov-o/#Generation

2 https://www.w3.org/TR/prov-o/#Revision

3 https://www.w3.org/TR/prov-o/#Invalidation

## References

1. International Society for Biocuration (2018) Biocuration: Distilling data into knowledge. PLoS Biol., 16, e2002846.

2. Tang, Y. A., Pichler, K., Füllgrabe, A., et al. (2019) Ten quick tips for biocuration. PLoS Comput. Biol., 15, e1006906.

3. Poux, S., Arighi, C. N., Magrane, M., et al. (2017) On expert curation and scalability: UniProtKB/Swiss-Prot as a case study. Bioinformatics, 33, 3454–3460.

4. Venkatesan, A., Karamanis, N., Ide-Smith, M., et al. (2019) Understanding life sciences data curation practices via user research. F1000Res., 8, 1622.

5. Holinski, A., Burke, M. L., Morgan, S. L., et al. (2020) Biocuration - mapping resources and needs. F1000Res., 9.

6. Bateman, A. (2010) Curators of the world unite: the International Society of Biocuration. Bioinformatics, 26, 991.

7. Attwood, T. K., Bongcam-Rudloff, E., Brazas, M. E., et al. (2015) Correction: GOBLET: The Global Organisation for Bioinformatics Learning, Education and Training. PLoS Comput. Biol., 11, e1004281.

8. Corpas, M., Jimenez, R. C., Bongcam-Rudloff, E., et al. (2015) The GOBLET training portal: a global repository of bioinformatics training materials, courses and trainers. Bioinformatics, 31, 140–142.

9. Rigden, D. J. and Fernández, X. M. (2020) The 27th annual Nucleic Acids Research database issue and molecular biology database collection. Nucleic Acids Res., 48, D1–D8.

10. Attwood, T. K., Agit, B. and Ellis, L. B. M. (2015) Longevity of Biological Databases. EMBnet.journal, 21, 803.

11. Perfetto, L., Pastrello, C., Del-Toro, N., et al. (2020) The IMEx coronavirus interactome: an evolving map of Coronaviridae-host molecular interactions. Database, 2020.

12. Panni, S., Lovering, R. C., Porras, P., et al. (2019) Non-coding RNA regulatory networks. Biochim. Biophys. Acta Gene Regul. Mech., 1863, 194417.

13. Haenig, C., Atias, N., Taylor, A. K., et al. (2020) Interactome Mapping Provides a Network of Neurodegenerative Disease Proteins and Uncovers Widespread Protein Aggregation in Affected Brains. Cell Rep., 32, 108050.

14. El-Gebali, S., Mistry, J., Bateman, A., et al. (2019) The Pfam protein families database in 2019. Nucleic Acids Res., 47, D427–D432.

15. Park, Y. M., Squizzato, S., Buso, N., et al. (2017) The EBI search engine: EBI search as a service-making biological data accessible for all. Nucleic Acids Res., 45, W545–W549.

16. Jassal, B., Matthews, L., Viteri, G., et al. (2020) The reactome pathway knowledgebase. Nucleic Acids Res., 48, D498–D503.

17. Haak, L. L., Fenner, M., Paglione, L., et al. (2012) ORCID: a system to uniquely identify researchers. Learn. Publ., 25, 259–264.

18. Griffith, M., Spies, N. C., Krysiak, K., et al. (2017) CIViC is a community knowledgebase for expert crowdsourcing the clinical interpretation of variants in cancer. Nat. Genet., 49, 170–174.

19. Hatos, A., Hajdu-Soltész, B., Monzon, A. M., et al. (2020) DisProt: intrinsic protein disorder annotation in 2020. Nucleic Acids Res., 48, D269–D276.

20. Huotari, K. and Hamari, J. (2012) Defining gamification: a service marketing perspective. Proceeding of the 16th International Academic MindTrek Conference, MindTrek ‘12, Association for Computing Machinery, New York, NY, USA, pp. 17–22.

21. Hamari, J., Koivisto, J. and Sarsa, H. (2014) Does Gamification Work? – A Literature Review of Empirical Studies on Gamification. In: 47th Hawaii International Conference on System Sciences. IEEE Computer Society Conference Publishing Services (CPS), p. 3025–3034.

22. Thessen, A.E., Woodburn, M., Koureas, D.et al. (2019) Proper Attribution for Curation and Maintenance of Research Collections: Metadata Recommendations of the RDA/TDWG Working Group. Proper Attribution for Curation and Maintenance of Research Collections: Metadata Recommendations of the RDA/TDWG Working Group.Data Sci. J.,18, 54.

23. Wimalaratne, S. M., Juty, N., Kunze, J., et al. (2018) Uniform resolution of compact identifiers for biomedical data. Sci Data, 5, 180029.

24. Moreau, L., Groth, P., Cheney, J., et al. (2015) The rationale of PROV. Journal of Web Semantics, 35, 235–257.

